# FLIM-FRET Imaging of AMPA Receptors: New Principle for Subtype-Specific Elucidation

**DOI:** 10.64898/2025.12.17.694835

**Authors:** Parisa Rezaee, Miriam G. Pedersen, Linda G. Zachariassen, Markus Staudt, Matthias M. Herth, Anders S. Kristensen

**Affiliations:** Department of Drug Design and Pharmacology, Faculty of Health and Medical Sciences, University of Copenhagen, Denmark

## Abstract

α-amino-3-hydroxy-5-methyl-4-isoxazolepropionic acid receptors (AMPARs) mediate fast excitatory neurotransmission, and their subunit composition (GluA1-4) critically shapes synaptic strength and plasticity. Discriminating AMPAR subtypes at the molecular level remains challenging with conventional techniques. Here, we applied Förster resonance energy transfer (FRET) combined with fluorescence-lifetime imaging microscopy (FLIM) to resolve subtype-specific AMPAR assemblies. Cyan fluorescent protein (CFP) and HALO domain were genetically introduced into GluA1–3 subunits to make intrareceptor FRET pairs. Self-labeling protein (SLP) domains such as SNAP and HALO were used with cell-impermeable substrates to selectively label surface-expressed receptors. This approach identified di-heterotetrameric GluA1/2 and GluA2/3 assemblies at the HEK293T cell membrane, whereas GluA1/3 failed to form di-heteromers. These findings demonstrate the FLIM-FRET method as a tool for differentiating AMPAR subtypes in living cells, providing a foundation for studying subtype-specific receptor organization in excitatory synapses.

## Introduction

α-amino-3-hydroxy-5-methyl-4-isoxazolepropionic acid receptors (AMPARs) are found in the presynaptic and postsynaptic membranes of the neural cells (1). Like other major subtypes of ionotropic glutamatergic receptors (iGluRs), such as N-methyl-D-aspartate (NMDA) and kainate receptors, AMPARs are ligand-gated ion channels. These receptors are tetrameric assemblies composed of four subunits, GluA1-4, which can form both homo- and heterotetramers that determine channel properties and synaptic function (2). Advances in X-ray crystallography and cryo-electron microscopy (cryo-EM) have provided structures of AMPARs in different functional states (3–7).

The four AMPAR subunits show both overlapping and distinct properties in terms of function, expression, and post-transcriptional regulation (14–16). The GluA1 subunit plays a pivotal role in activity-dependent synaptic plasticity essential for learning and memory (8,9). The GluA2 subunit is the key regulator of calcium ion impermeability in AMPARs, which is crucial for neuronal survival and maintaining synaptic stability (1,10). The glutamine/arginine (Q/R) site, which is found only in GluA2 influences single-channel conductance, polyamine block, and subunit assembly into heteromeric AMPARs (11). GluA1 preferentially forms heterotetramers with GluA2, but can also assemble as homotetramers (12). AMPAR subtypes containing GluA3 predominantly form as heterotetramers with GluA2, rather than homotetramers of GluA3 (13). Mechanisms for regulation of receptor function are subtype-specific (1,16). GluA1 is modulated by phosphorylation at serines 845 and 831, which controls receptor trafficking and synaptic retention (17). GluA2 regulation occurs through RNA editing at the Q/R site. The unedited form with glutamine tends to be retained in the endoplasmic reticulum and undergoes less efficient trafficking to the plasma membrane; in contrast, the edited form with arginine efficiently traffics to synapses and promotes stable surface expression of AMPARs (18). GluA3 exhibits slower trafficking from the endoplasmic reticulum, which is governed by its intracellular C-terminal domains and interactions with auxiliary subunits, such as TARPs and scaffold proteins, that direct synaptic targeting and stability (19,20).

Subunit arrangement and dimensions of AMPARs are known from determined X-ray and cryo-EM structures (4–6,21). These predicted distances between subunits are perfectly suited for the use of a biophysical tool such as Förster resonance energy transfer (FRET) to detect if two different subunits are present in the same AMPAR subtype. FRET, combined with fluorescence lifetime imaging microscopy (FLIM), an imaging method that measures changes in fluorescence decay times sensitive to molecular proximity, can be applied to determine whether two different AMPAR subunits are assembled in the same receptor complex. This method enables the detection of receptor subtypes expressing specific subunit combinations in live cells or tissue samples.

Discriminating among AMPAR subtypes is crucial for understanding their distinct roles in physiological processes and for developing targeted therapeutic interventions. Physiologically, Ca^2+^-permeable (CP) and Ca^2+^-impermeable (CI) AMPARs differ significantly in their ion conductance properties, influencing synaptic plasticity, neuronal excitability, and calcium signaling pathways that underlie learning, memory, and neuronal survival (1). Therapeutically, CP-AMPARs contribute to excitotoxicity in pathological conditions such as ischemia, epilepsy, and neurodegeneration, making them specific targets for drug development aimed at reducing calcium-mediated neuronal damage (22). Therefore, identifying the presence and distribution of AMPAR subtypes with different Ca^2+^ permeability is essential for both basic neuroscience research and clinical approaches aimed at modulating synaptic function and protecting neurons (10). The inability to resolve receptor subtype composition at individual synapses limits progress in understanding the precise physiological roles of AMPARs and how they contribute to synaptic plasticity and neuronal signaling. Synaptic responses can differ based on whether GluA1/2 or GluA2/3 receptor subtypes predominate, affecting calcium permeability, receptor kinetics, and plasticity mechanisms (23). This lack of resolution hinders dissecting how specific AMPAR subtypes contribute to learning, memory, and pathological states (24,25).

In this study, we calculated FRET efficiency (E_FRET_) within the expected range, where the distance between donor and acceptor is less than 100 Å for cells expressing GluA1 and GluA2 subunits and GluA2 and GluA3 subunits. The calculated distance demonstrates that FLIM-FRET can be applied to differentiate AMPAR subtypes, validating GluA1/2 and GluA2/3 assemblies. No FRET was detected between GluA1 and GluA3. The study thus provides methodological and complementary evidence for AMPAR subtype discrimination.

## Results

### Expression of fluorescent-enabled AMPARs

To express GluA1, GluA2(R), and GluA3 with a fluorescence marker in HEK293T cells, Cyan fluorescent protein (CFP) and HALO domain were inserted between the signal peptide (SP) and N-terminal domain (NTD) of plasmid DNAs encoding AMPARs (Fig.1.a). The pmLINK vector, a dual-expression vector with two expression cassettes and CAG promoters used to ensure equal transcription of the inserted DNA fragments (26). The GluA1, GluA2(R), and GluA3 were tagged with HALO domain and CFP, to generate three plasmid DNA constructs. To test if the plasmid DNA constructs expressed the AMPAR subunits with CFP and HALO domain, HEK293T cells were transfected with each plasmid DNA. The cells expressing CFPGluA1HALOGluA2(R) (CA1HA2(R)), CFPGluA2(R)HALOGluA3 (CA2(R)HA3), and CFPGluA3HALOGluA1 (CA3HA1) after labeling with HALO-Alexa Fluor 488 (AF488), showed fluorescence signals for both CFP and HALO domain (Data S1). The results from the imaging of the HEK293T cells transfected with the three plasmid DNA constructs showed GluA1, GluA2(R), and GluA3 expressed in the cell membrane.

**Figure 1.**
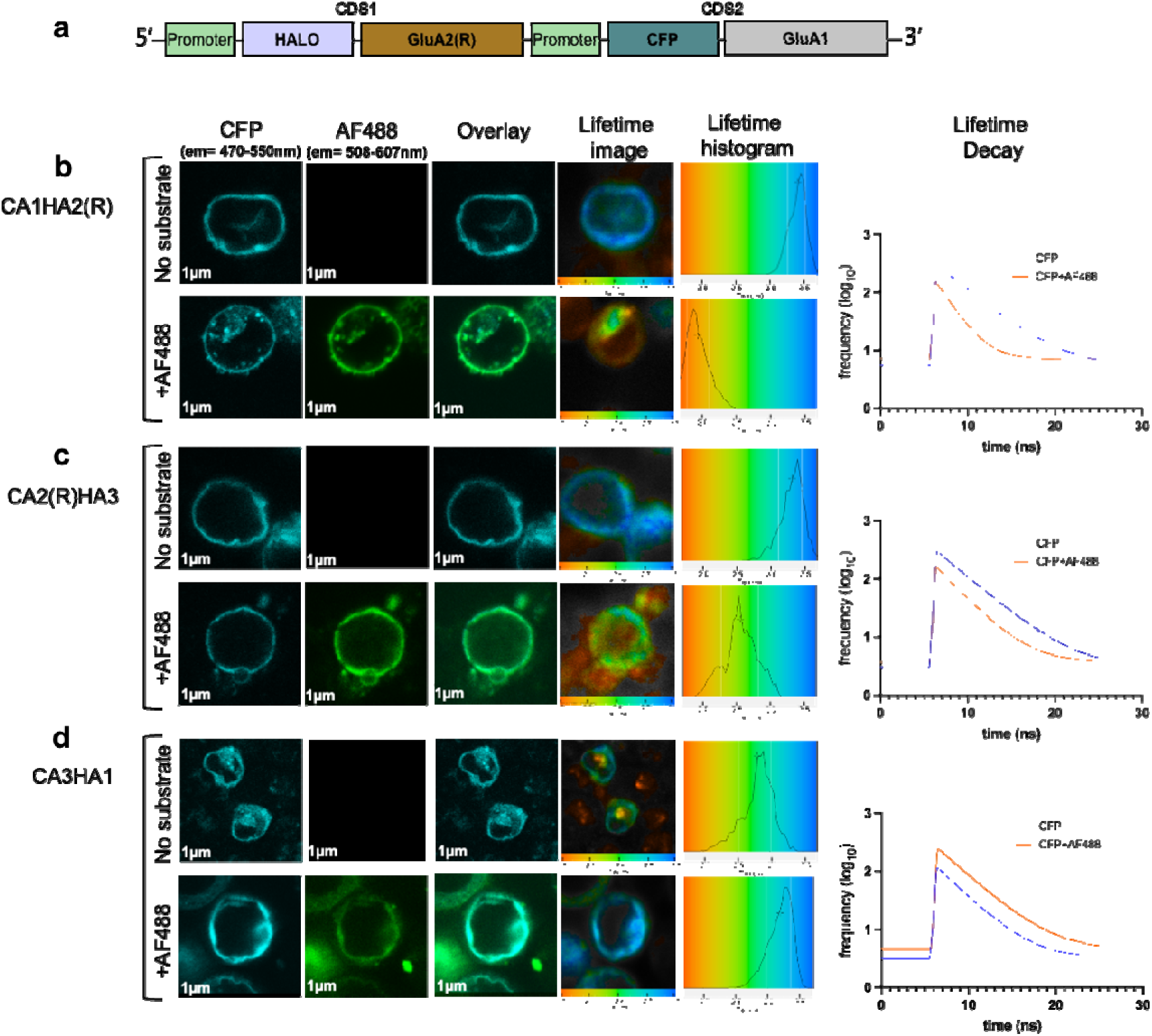
Design of FRET-enabled dual AMPARs expressing CFP and HALO and FRET evaluation. **(a)** Schematic overview of GluA1,2 subunits with CFP and HALO domain inserted. **(b,c)** The CFP lifetime images show that after adding acceptor, the lifetime shifts from blue to green-orange, which represents the decrease in donor lifetime in single cells expressing CA1HA2(R) and CA2(R)HA3. The lifetime histogram for the single cells expressing GluA1/2 and GluA2/3 in the presence of the acceptor is shifted to the left, which indicates a lower lifetime. Whereas, there is no decline in lifetime for cells expressing **(d)** CA3HA1. The CFP lifetime histogram for the single cells expressing GluA1, 3 after adding the acceptor does not represent a lower lifetime compared to CFP alone (FLIM and lifetime histograms color scale bar shows 1.7-3.7 ns from red to blue). **(b,c,d)** On the right, lifetime decay curves of the single cells expressing CA1HA2(R), CA2(R)HA3, and CA3HA1 are shown (CFP: blue, CFP/AF488: orange). The CFP lifetime decay curve in the blue color shows a mono-exponential decay curve in the absence of an acceptor. In the presence of AF488, the curve adopts a multi-exponential decay curve except for cells expressing CA3HA1.

### Characterization of FRET between AMPAR subunits

To determine FRET, CFP was chosen as the FRET donor. An optimal FRET donor shows single-exponential lifetime decay, and its emission spectrum overlaps with the acceptor absorption spectrum. The mCerulean3, a variant of CFP, with enhanced brightness, fulfills these criteria (27). HALO domain was labeled with a membrane-impermeable substrate, AF488. The CFP emission spectrum and AF488 excitation spectrum overlap by more than 75%, facilitating maximum fluorescence energy transfer from the donor to the acceptor. Since overlapping between the donor emission spectrum and acceptor excitation spectrum affects E_FRET_, CFP/AF488 was selected as the FRET pair. Inserting donor and acceptor between SP and NTD of subunits is expected to give donor/acceptor distance in di-heterotetrameric receptors around 40 to 60 Å (4,6,21) and display FRET. First, by selecting the single cells with the best membrane expression of CFP, the confocal and lifetime images of CFP for single cells expressing CA1HA2(R), CA2(R)HA3, and CA3HA1 were obtained in the absence of HALO substrate (Fig.1.b-d). Then, the fluorescence images of selected cells were taken and CFP lifetime was measured in the presence of AF488 (Fig.1.b-d).

CFP lifetime mean in the absence of acceptor for cells expressing CA1HA2(R), CA2(R)HA3, and CA3HA1 was 3.30±0.16, 3.44±0.20, and 2.94±0.12, respectively (Table 1). In cells expressing CA1HA2(R), the CFP lifetime mean significantly decreased to 2.03±0.10 in the presence of AF488 (Table 1). Figure 2b shows FLIM images for CFP using color coding that reflects the donor lifetime in the individual pixels for single cells expressing CA1HGA2(R) in the presence of AF488. The same color coding is used in histogram plots of the frequency of pixels lifetime. CFP lifetime images and lifetime decay histograms for single cells expressing with CA2(R)HA3 in the presence of AF488 indicate a significant decrease in the lifetime (Fig. 1.c) and CFP lifetime mean decrease to 2.51±0.06 (Table 1). In the cells expressing CA3HA1, there was no decline in CFP lifetime in the presence of AF488 (Table 1, Fig. 1.d). FLIM images and lifetime decay histograms align with this data (Fig. 1.d). The lifetime decay curve for cells expressing CA1HA2(R) shows a single-exponential decay for CFP without an acceptor. After labeling the HALO domain, the lifetime decay curve in the presence of AF488 is multi-exponential (Fig. 1.b). For cells expressing CA2(R)HA3, the CFP lifetime decay curve after labeling with AF488 shows multi-exponential lifetime decay (Fig. 1.c). In the cells expressing CA3HA1, the CFP lifetime decay shows the same decay curve fit before and after labeling the HALO domain with AF488 (Fig. 1.d).

**Figure 2.**
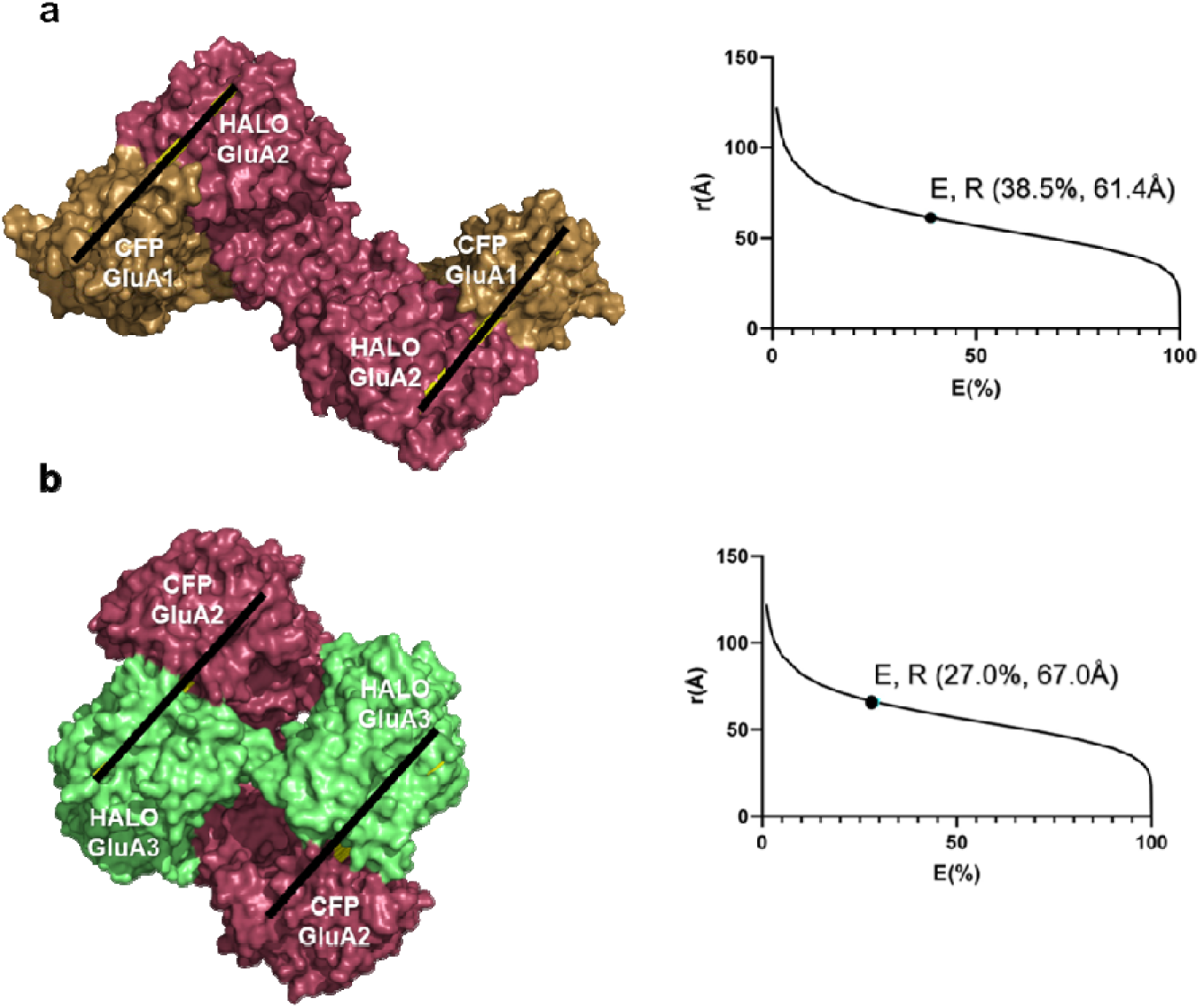
Surface view of GluA1/2 and GluA2/3 NTD with intersubtype distance representation. On the right shows the surface view of NTD for di-heterotetramer receptor **(a)** GluA1/2 (PDB:7OCC) and **(b)** GluA2/3 (PDB: 5FWY) (GluA1 subunit: sand color, GluA2 subunit: ruby color, GluA3 subunit: green color). The distance is measured approximate region where the CFP and HALO domain are inserted. **(a)** The representative distance on average is ∼61Å, the same as the calculated distance shown on the right using E_FRET_. In Figure **(b)**, the distance is ∼65Å, which is close to what we calculate in the graph based on FLIM-FRET measurements (Sample size is the number of cells E_FRET_ calculated from *(Table 1)*).

**Table 1.** Fluorescent lifetime and E_FRET_ of dual CFP/HALO tagged GluA1-3 receptors. Donor lifetime τ ± SEM of *n* cells obtained by FLIM-FRET measurements of dual CFP/HALO tagged GluA1-3 receptors expressed in HEK293T cells. The HALO domain was labelled with AF488. FLIM analysis was conducted as described in *Material and Method* E_FRET_ (%) calculated from equation 1.

| Construct | Donor | Acceptor | Donor lifetime $\tau$ (ns) | n | $E_{\text{FRET}}$ (%) |
| --- | --- | --- | --- | --- | --- |
| CA1HA2(R) | CFP | - | $3.30 \pm 0.16$ | 6 | - |
| | CFP | AF488 | $2.03 \pm 0.10$ | 12 | 38.48 |
| CA2(R)HA3 | CFP | - | $3.44 \pm 0.20$ | 9 | - |
| | CFP | AF488 | $2.51 \pm 0.06$ | 3 | 27.03 |
| CA3HA1 | CFP | - | $2.94 \pm 0.12$ | 3 | - |
| | CFP | AF488 | $3.23 \pm 0.07$ | 5 | No FRET |

CFP/AF488 pair showed E_FRET_ of 38.48% and 27.03% for cells expressing CA1HA2(R) and CA2(R)HA3, respectively (Table 1). The decrease in the CFP lifetime and the calculated E_FRET_ in the presence of an acceptor in cells expressing CA1HA2(R) and CA2(R)HA3 showed that GluA1 and GluA2(R) subunits assemble as GluA1/2 subtype and GluA2(R) and GluA3 subunits as GluA2/3 di-heterotetramer subtype. Whereas GluA1 and GluA3 do not form a heterotetrameric subtype. Constructs containing GluA2(R), specifically GluA1/2 and GluA2/3, demonstrated strong expression. This outcome is consistent with the well-established role of GluA2 in promoting efficient AMPAR assembly and ER export, even in the absence of auxiliary subunits (28,29). In contrast, the GluA1,3 combination lacks the stabilizing influence of GluA2. As a result, proper folding, assembly, trafficking, and surface expression of GluA1/3 in HEK293T cells depend on co-expression with TARPs or other auxiliary subunits (30).

### Differentiating GluA1/2 from GluA2/3 subtype

To discriminate AMPAR GluA1/2 from GluA2/3, the distance between donor and acceptor was calculated. The calculated E_FRET_ from cells expressing CA1HA2(R) and CA2(R)HA3, labeled with AF488 was placed in eq2 (Material and method). The distance between donor and acceptor in the GluA1/2 subtype was 61.4Å, while the distance between the FRET pair in the GluA2/3 subtype was 67Å (Fig.2). In a study using cryo-EM showed the distance between GluA1 and GluA2 in heteromeric GluA1/2 AMPAR is approximately 60Å this data is in accordance with the calculated distance between donor and acceptor in GluA1/2 subtype (Fig.2a)(20). The distance between subunits of GluA2/3 in resting state, according to cryo-EM is ∼68Å (21), which is consistent with the calculated distance between donor and acceptor in cells expressing CA2(R)HA3 (Fig.2b). Using NTD-GluA1/2 structure (PDB: 7OCC) to depict the distance between donor and acceptor, the average calculated distance was ∼61Å (Fig.2a). The calculated distance between donor and acceptor is shown on the crystal structure of GluA2/3 NTD (PDB: 5FWY) (Fig.2.b). The calculated distance between donor and acceptor for subtypes GluA1/2 and GluA2/3 is an indicator to differentiate these two subtypes from each other.

The measured distance using the FLIM-FRET method is in accordance with AMPAR cryo-EM studies. According to the measured distance between donor and acceptor in GluA1/2 and GluA2/3, we indicated that the subunits GluA1 and GluA2(R) are 6.4Å closer compared to the distance between GluA2(R) and GluA3 subunits.

### Expression of self-labeling protein (SLP)-enabled dual AMPARs

To selectively collect fluorescence signals from AMPARs in the cell membrane, CFP was replaced with SNAP, a SLP domain (Fig.3.a). Biorthogonal reaction of cell impermeable ligands with genetically inserted SLP domains indicated fluorescence signals and FRET from the AMPARs on the cell membrane. To test if the pmLINK constructs expressed the AMPAR subunits with SNAP and HALO domains, HEK293T cells were transfected with plasmid DNA constructs. HEK293T cells expressing SNAPGluA1HALOGluA2(R) (SA1HA2(R)), HA1SA2(R), HALOGluA2(R)SNAPGluA3 (HA2(R)SA3), and SNAPGluA3HALOGluA1 (SA3HA1), after labeling with SNAP-AF488 and HALO-AF568 individually, showed fluorescence signals for both SNAP and HALO domains (Data S2).

**Figure 3.**
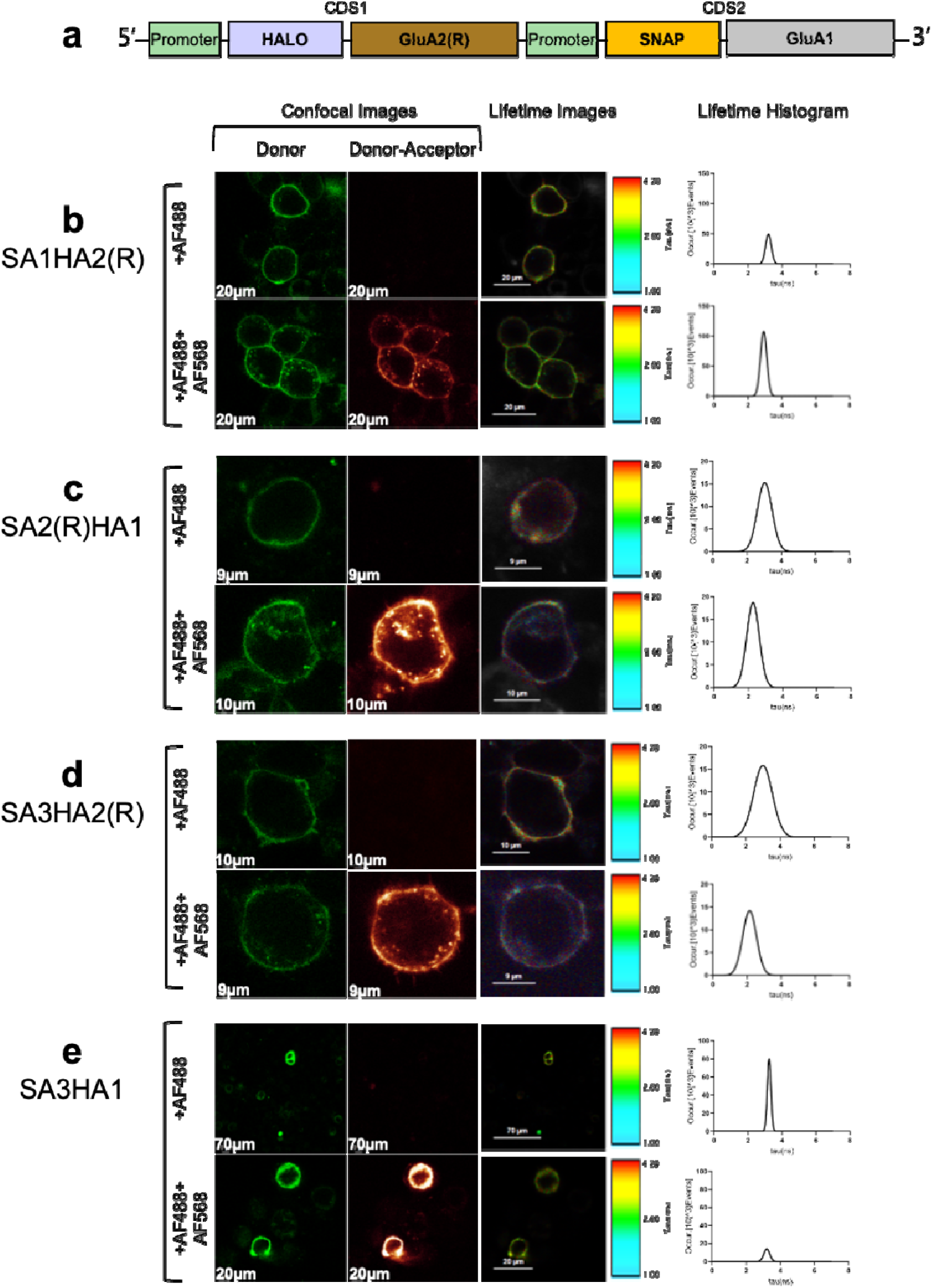
Design of FRET-enabled dual AMPARs expressing SNAP and HALO domains and FRET evaluation. **(a)** Schematic overview of GluA1,2 subunits with SNAP and HALO domains inserted. **(b-e)** Confocal images of fluorescence single cells expressing the SA1HA2(R), SA2(R)HA1, SA3HA2(R), and SA3HA1 constructs obtained by confocal laser scanning microscopy. The fluorescence lifetime of SNAP-AF488 in individual pixels was obtained by TCSPC analysis *(Material and Method)* and used to construct fluorescence lifetime images. In figures **(b, c)**, cells expressing GluA1,2 and **(d)** GluA2,3 labeled with SNAP-AF488 as FRET donor show a shift from a higher lifetime to a lower lifetime in the lifetime images and histograms in the presence of the acceptors HALO-AF568. **(e)** After the expression of GluA1,3 and adding the acceptor, the peak of lifetime in the histogram does not shift to a lower lifetime. (The FLIM color scale bar shows 1.00-4.20 ns from blue to red).

### Characterization of FRET between dual SLP-AMPAR subunits

FLIM-based FRET measurements in cells expressing fluorescently labeled GluA1–3 subunits enabled quantitative determination of the membrane distribution and relative abundance of distinct AMPAR subtypes. On the day of FLIM measurement, cells expressing SA1HA2(R), SA2(R)HA1, SA3HA2(R), and SA3HA1 were labeled with SNAP-AF488 and HALO-AF568 (Material and Method). By selecting the single cells with the best membrane expression of the SNAP domain, the confocal and lifetime images of SNAP-AF488 were obtained in the absence of the HALO substrates (Fig.3). Then, the selected cells were imaged, and SNAP-AF488 lifetime was measured in the presence of HALO-AF568.

SNAP-AF488 lifetime mean in the absence of acceptor for cells expressing SA1HA2(R), SA2(R)HA1, SA3HA2(R), and SA3HA1, respectively, was 3.27±0.16 ns, 3.11±0.21 ns, 3.03±0.16 ns, and 3.07±0.28 ns (Table 2). In the cells transfected with pmLINK-S-A1-H-A2(R), the SNAP-AF488 lifetime mean changed to 2.94±0.08 ns in the presence of AF568. In cells expressing SA2(R)HA1, the donor lifetime mean declined to 2.31±0.20 ns in the presence of AF568. In cells expressing SA3HA2(R), the donor lifetime mean declined to 2.50±0.16 ns in the presence of AF568 (Table 2).

**Table 2.**
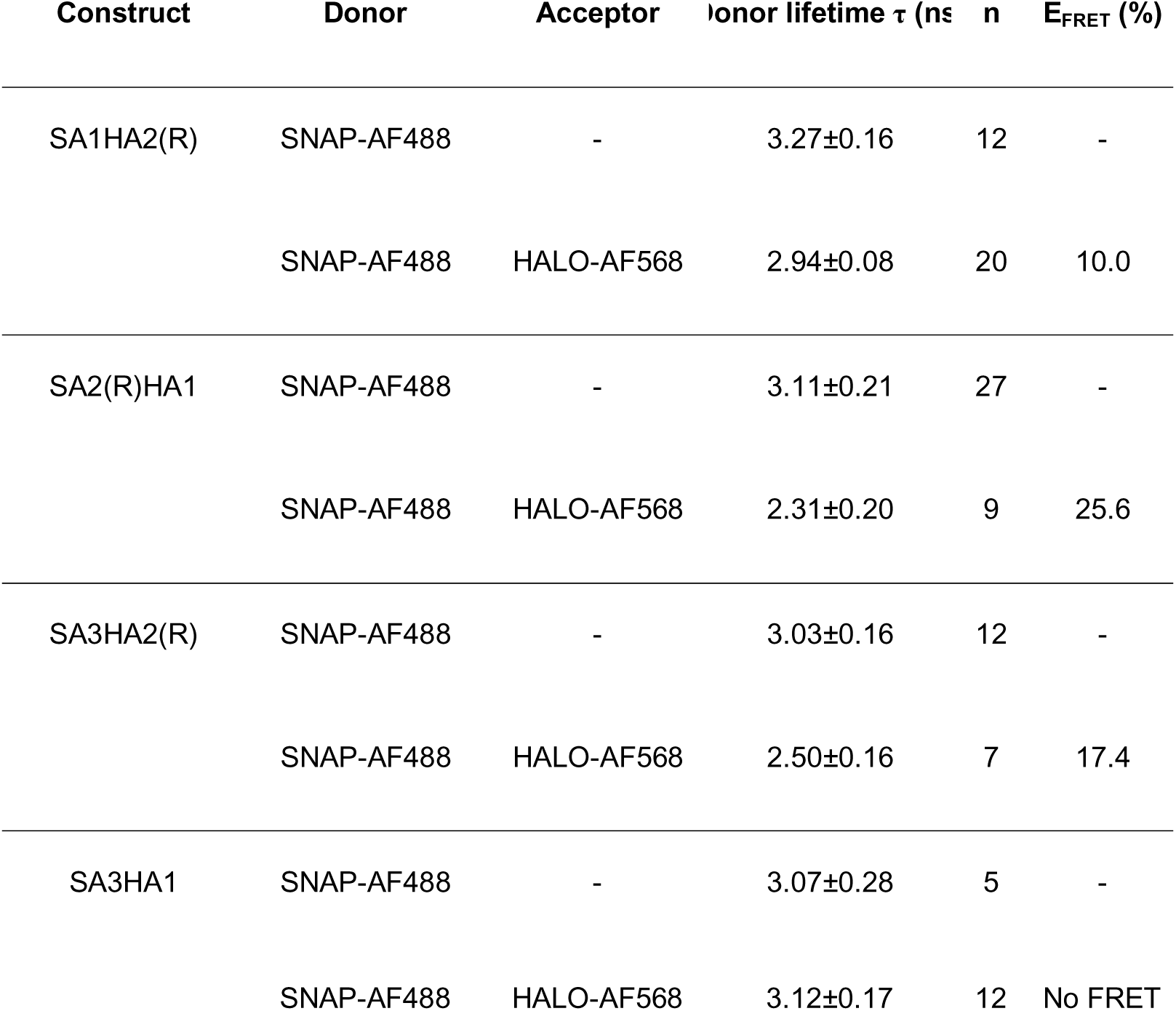
Fluorescent lifetime and E_FRET_ of dual SNAP/HALO tagged GluA1-3 receptors. Donor lifetime τ ± SEM of n cells obtained by FLIM-FRET measurements of SNAP/HALO tagged GluA1-3 receptors expressed in HEK293T cells. SNAP and HALO domains labelled with SNAP-AF488 and HALO-AF568.

For GluA1,3 combination, the FLIM measurements showed there is no decline in the lifetime of the donor in the presence of the acceptor. SNAP-AF488 lifetime images and lifetime decay histograms for single cells expressing SA1HA2(R), SA2(R)HA1, SA3HA2(R), and SA3HA1 align with this data. The significant change in the lifetime image and lifetime decay histogram of SNAP-AF488 in the presence of AF568 indicates a shift to a lower lifetime (Fig. 3.b-d). Whereas in cells expressing SA3HA1, following the calculated lifetime mean of SNAP-AF488, there was no detectable decline in the donor lifetime in the presence of the acceptor. The lifetime image and lifetime decay histogram of the single cell did not indicate a shift to a lower lifetime (Fig.3.e). The presence of the FRET in cells expressing SA1HA2(R), SA2(R)HA1, and SA3HA2(R), showed that GluA1 and GluA2(R), as well as GluA2(R) and GluA3, are di-heterotetramers on the membrane. The absence of the FRET in cells with expressing SA3HA1 showed that GluA1 and GluA3 do not represent di-heterotetrameric subtypes on the membrane and appear as a combination of GluA1 homotetramers and GluA3 homotetramers.

To distinguish AMPAR GluA1/2 from GluA2/3, the distance between subunits within each subtype was calculated. The calculated E_FRET_ in cells expressing SA1HA2(R), SA2(R)HA1, and SA3HA2(R), was placed in eq2 (Material and Method). The FRET-derived distance between GluA1 and GluA2 depends on the insertion site of the fluorescence domain and the adjacent subunits. As a result, swapping HALO and SNAP between GluA1 and GluA2 changed the calculated distance. The calculated distance between subunits with donor and acceptor in cells expressing SA1HA2(R) was 81.90Å, whereas in cells expressing SA2(R)HA1, the calculated distance was 67.80Å (Fig. 4. a,b). The distance between subunits GluA2(R) and GluA3 in cells expressing SA3HA2(R) was calculated to be 73.62Å (Fig. 4.c).

**Figure 4.**
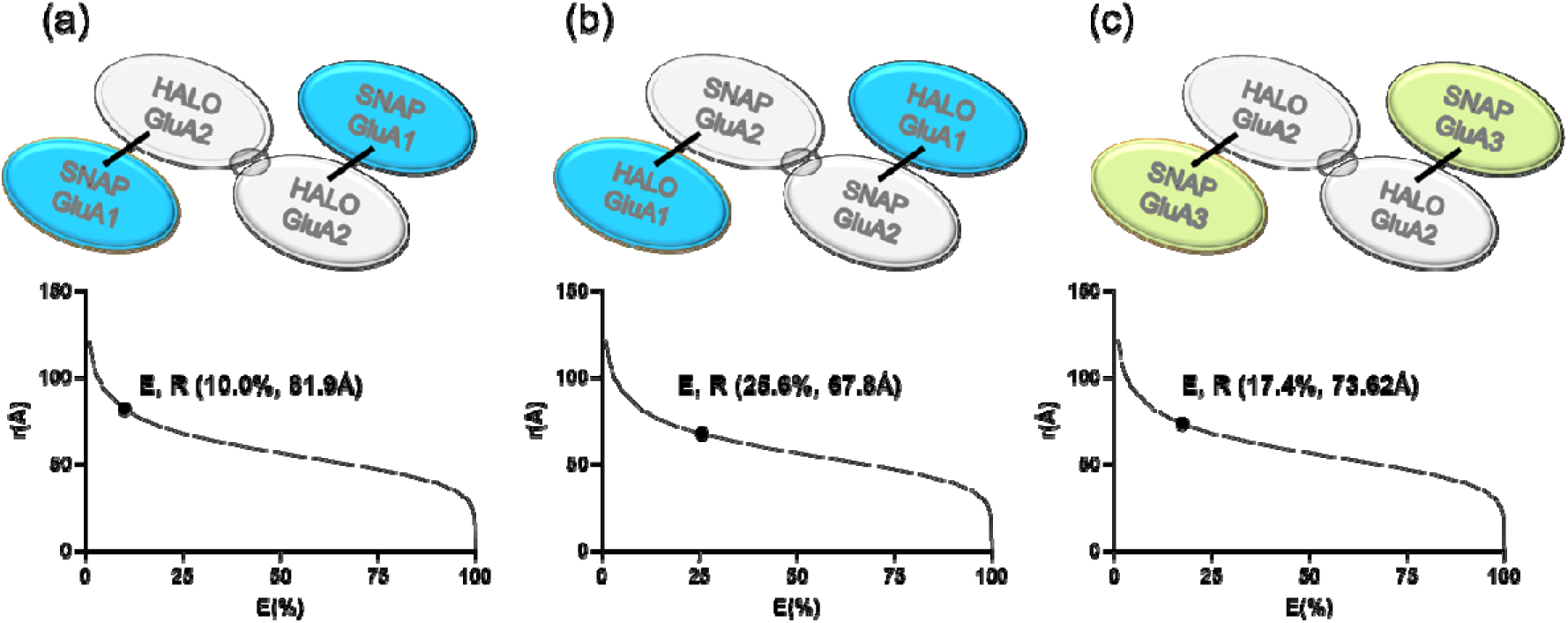
Calculated distances between donor and acceptor in AMPAR subtypes SA1HA2(R), SA2(R)HA1, and SA3HA2(R) The distance between SNAP domain labeled with AF488 and HALO domain labeled with AF568 was calculated by using E_FRET_. In the schematic view GluA1 subunit is in grey color, GluA2 subunit is in blue color, and GluA3 subunit is in green color. The distance between subunits of each AMPAR complex is less than 100Å and is shown as a black line from one subunit to another. **(a)** Approximate distance between HALOGluA2 and SNAPGluA1 is 81.9Å, whereas **(b)** the distance between SNAPGluA2 and HALOGluA1 is 67.8Å. **(c)** The distance between HALOGluA2 and SNAPGluA3 is 73.62Å (Sample size is the number of cells, fluorescence lifetime mean was calculated and placed in eq 1 *(Material and Method, Table 2)*).

### Effect of Fixation on FRET Measurements in HEK293T Cells

To assess whether fixation of HEK293T cells impacts E_FRET_, the pmLINK-S-A2(R)-H-A1 DNA construct was selected as a representative di-heterotetrameric AMPAR subtype, due to its robust expression and established E_FRET_ in prior measurements. HEK293T cells were transfected and labeled with SNAP-AF488 and HALO-AF568 according to the same procedure. Cells fixated by a chemical-based assay, using formaldehyde (Material and Methods) was advantageous due to its compatibility with the FRET technique and preserving cellular morphology and structure (33). FLIM was recorded before fixation and at 1-, 7-, and 21-day post-fixation. The FLIM recording indicated that SNAP-AF488 lifetime decreased due to fixation of HEK293Tcells for cells labeled with donor alone and cells labeled with donor/acceptor (Fig. 5). In seven days post-fixation, E_FRET_ for SNAP-AF488/HALO-AF568 decreased compared to one day post-fixation. In 21-day fixed cells, the E_FRET_ for SNAP-AF488/HALO-AF568 was very low (Fig. 5). These results demonstrate that FRET remains detectable in HEK293T cells up to 7 days post-fixation. Notably, E_FRET_ was significantly higher at 1 day post-fixation for the AF568 pair and remained measurable after 7 days, highlighting the stability of this fluorophore pairing under fixed conditions.

**Figure 5.**
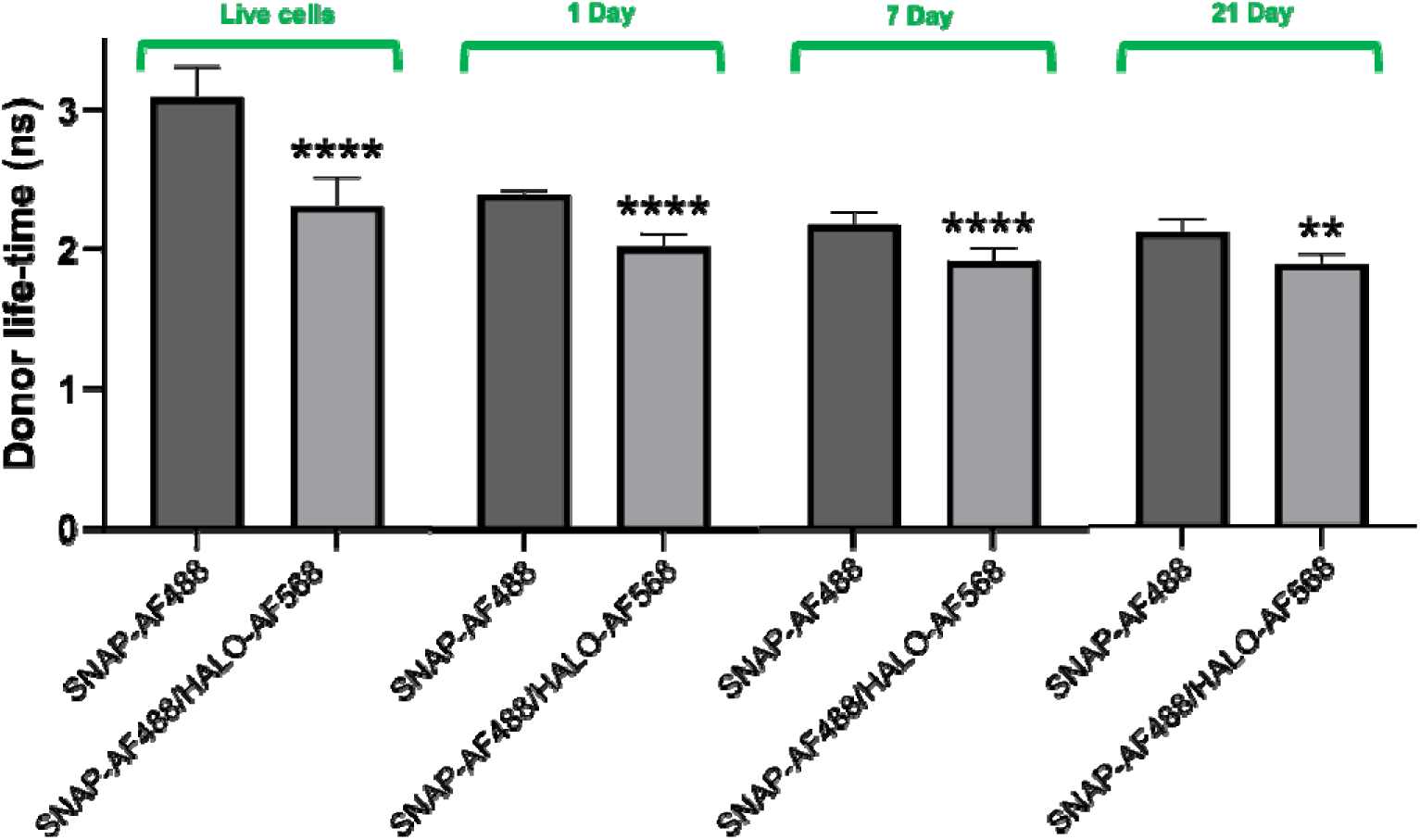
Effect of HEK293T cells fixation on FRET. Column-bar histogram of transfected cells with pmLINK-S-A2(R)-H-A1 after labelling with SNAP and HALO ligands indicated a drop in SNAP-AF488 in the absence and presence of HALO-AF568. In 1 and 7 days post-fixation SNAP-AF488/HALO-AF568 indicates FRET. Statistical analysis by ordinary one-way ANOVA test, followed by Dunnett’s post hoc test, indicates (**** p<0.0001. ** p=0,0035 (N_cells_ = 9-22 (live cells), 6-9 (1-day post-fixation), 7-9 (7 days post-fixation), 4-5 (21 days post-fixation)).

### Discussion

The subunit combination of AMPAR determines essential receptor characteristics such as Ca^2+^ ion permeation and gating kinetics, which in turn influence synaptic physiology (34,35). Differentiating CP-AMPARs from CI-AMPARs is highly demanded due to the involvement of overexpressed CP-AMPARs in excitotoxity, neural injury, epilepsy, and chronic pain (36,37). AMPARs not only play a crucial role in initiating excitatory signaling but also modulate synaptic strength through dynamic trafficking at the postsynaptic site (38–40). This process is fundamental to different types of experience-dependent synaptic plasticity (41). A study of AMPAR composition indicated that GluA1/2 and GluA2/3 are dominant AMPAR heteromers in hippocampal CA1 pyramidal neurons (23). AU-FDS (13) and cryo-EM (31,42) have provided information about the subunit arrangement and preferences of AMPAR compositions, as well as the heteromeric structures of GluA1/2 and GluA2/3 subtypes. However, available information on how GluA1 and GluA3 assemble and the abundance of GluA1/3 composition is limited. Moreover, there are few established methods to distinguish between different heterotetrameric AMPAR subtypes.

To discriminate AMPAR subtypes, we used FLIM technique to measure the lifetime variation of the donor in the presence of an acceptor to calculate the E_FRET_. The distance between donor and acceptor in this study indicates that GluA1,2 and GluA2,3 subunits formed GluA1/2 and GluA2/3 di-heterotetramer complexes, respectively. No FRET was detected for cells transfected with GluA1,3 subunits, indicating that the distance between the subunit with the donor and the subunit with the acceptor is more than 100Å.

In a study focused on CA1/CA2 pyramidal neurons, using immunoprecipitation and limited antibodies only selective for GluA1 and GluA2/3, it has been shown that major AMPAR complexes are composed of GluA1,2 or GluA2,3, while GluA1,3 was the minor complex (43). Applying immunoprecipitation and using anti-GluA2/3 antibodies eliminated all GluA3 immunoreactivity in the depleted fraction. In contrast, immunoprecipitation with anti-GluA1 antibodies retained approximately 90% of the GluA3 in the depleted fraction. These findings indicated that a limited subset of receptor complexes includes both GluA1 and GluA3, while the majority of complexes consist of either GluA1 and GluA2 or GluA3 and GluA2. Immunoreactivity for all subunits was observed in the bound fraction, except for GluA3, which exhibited only a negligible band. This finding aligns with the analysis of the unbound fraction, indicating that antibodies targeting GluA1 do not significantly co-immunoprecipitate GluA3 (43). The study by Wenthold et al., 1996 indicates the major and minor AMPARs complexes combination, but does not provide direct information about homo/heterotetrameric combination of AMPAR due to limited knowledge and approaches to study the receptor composition. The results we presented here regarding the presence of FRET for GluA1/2 and GluA2/3 are consistent with the conclusion from Wenthold et al., 1996. They indicated 70% of GluA2 is associated with GluA1, and anti-GluA2/3 antibody removed all GluA2/3 immunoreactivity. Furthermore, they indicated 90% of GluA3 does not immunoprecipitate with GluA1, which indirectly relates to the results in our study regarding undetectable FRET for GluA1/3 subtype. It is important to consider that the Wenthold et al., 1996 study is based on the early assumption that AMPAR is a pentameric receptor complex.

A recent study suggests that the exposed dimer interfaces in the LBD of GluA3(R) facilitate its assembly into heteromeric AMPAR complexes (19). This was demonstrated by patch-clamp recordings in HEK293 cells, where even a fourfold excess of GluA3(R) over GluA2(R) predominantly resulted in non-rectifying currents, indicative of GluA2(R)/GluA3(R) heteromers. In contrast, under identical conditions, GluA3(G) and GluA1 mainly formed homomers, as evidenced by their strongly rectifying current profiles (19). These findings from Pokharna et al., 2025 align with earlier studies showing that the mammalian GluA3(R) isoform tends to be retained in the ER, unless it either forms heteromeric assemblies or undergoes mutation at position 439 in the LBD, where arginine is substituted with glycine (21,44,45). In this study, because we used the G isoform of GluA3 in co-expression experiments with GluA1 and GluA2 to assemble GluA3-containing heterotetramers, it is worth considering a future modification: expressing GluA3(R) together with GluA1, to promote GluA1/3(R) heteromer formation.

We indicated the expression of GluA1/2 and GluA2/3 in HEK293T cells. Hence, the technique applied in this work does not limit the detection of GluA1/3. The first conclusion is that the GluA1/3 complex in the HEK293T cells transfected with GluA1 and GluA3 is not formed. We report that constructs containing GluA2(R) (GluA1/2, GluA2/3) show strong expression because GluA2 promotes efficient assembly and ER export of AMPARs even without TARPs (28,29). In contrast, GluA1/3 lacks GluA2-mediated stabilization and therefore would require TARP co-expression for proper folding, assembly, surface expression, and trafficking (29,30). The second conclusion is that transfected HEK293T cells expressing CP-AMPARs, GluA1,3, underwent excitotoxicity. The excitotoxicity led to a high number of dead cells and difficulty in FLIM recording.

Overall, this work contributes insights into differentiating AMPAR di-di-heterotetrameric complexes, highlighting the complexities of AMPAR composition and the methodological challenges in detecting AMPAR subtypes. The findings underscore the need for refined techniques to capture the diversity of AMPAR compositions and their implications for synaptic function.

## Acknowledgment

We thank Kasper B. Hansen for helpful discussion.

## Material and Method

**Materials:** Chemicals and reagents are described in detail in SI.

### Molecular Biology

#### Plasmid DNA constructs

The mammalian expression plasmid pmLINK was used as a vector for all FRET-enabled AMPAR constructs. Modified versions of the rat GluA1 and GluA3 DNA sequences with EcorV restriction site and GluA2 with AgeI restriction site in NTD were designed for insertion of CFP/SLP via Vector NTI software (Invitrogen, Carlsbad, USA). To prepare pmLINK-C-A1, pmLINK-C-A2, and pmLINK-C-A3 fusion constructs, first, CFP in the form of the CFP-variant mCerulean3 (27) cassette was amplified by using primers (SI) for PCR amplification. Then, CFP was inserted into the plasmid DNA coding sequence of GluA1, GluA2, and GluA3 in restriction-digested pmLINK using directional cloning of DNA fragments by use of the In-Fusion cloning kit, HiFi master mix (Invitrogen), according to the manufacturer’s protocol. The DNA constructs, pmLINK-H-A1, pmLINK-H-A2(R), and pmLINK-H-GluA3, were prepared. To construct dual pmLINK DNA constructs (Fig. 1.a), first CFP and HALO domains were inserted between SP and NTD of the subunits GluA1, GluA2(R), and GluA3 via an in-fusion cloning technique. pmLINK vector has one upstream (LINK1) and downstream (LINK2) of the expression CAG promoters (26). LINK1 contains a PacI restriction enzyme site, and LINK2 has a SwaI and PacI recognition site.

Second, pmLINK-H-A1, pmLINK-H-A2(R), and pmLINK-H-A3, each digested with a SwaI restriction enzyme. The insert DNA fragments, pmLINK-C-A1, pmLINK-C-A2(R), and pmLINK-C-A3 were digested at two PacI restriction sites. Plasmid DNAs were incubated with restriction enzymes for 14 hours at 37°C. Digested DNAs with PacI resulted in two DNA fragments. The DNA fragments with CFP and receptor sequences were separated and purified. After purification of all vector DNAs, a NEBuilder HiFi DNA Assembly Master Mix kit was used (New England Biolabs, USA) at 50°C for 30 minutes according to the manufacturer’s recommendation to fuse linearized vectors.

To prepare fluorescent-enabled dual AMPARs with SNAP and HALO domains, four pmLINK-plasmid DNA constructs were prepared (Fig. 2.a) in the same procedure as described for dual AMPARs containing CFP and HALO domain was repeated.

### Cellular Biology

#### HEK293T Cell culture and transfection

HEK293T cells from ATCC were cultured in DMEM supplemented with 10% FBS, 100 µg/ml penicillin, and 100 units/ml P/S. On the first day, cells were seeded on the black 96-well glass plate, 0.32 cm^2^, previously coated with PDL. After 24 hours, cells reached 40-60% confluency and were transfected with 0.1µg/well plasmid DNA using LipoD293 reagent (DNA: LipoD293: DMEM ratio of 1:3:30). After 48 hours, cells were labeled in individual wells with 2 µM of HALO-AF488 and HALO-AF568 for 45 minutes at 37°C. Cells were washed twice with Phosphate-Buffered Saline with Calcium and Magnesium (PBSCM: 250µl MgCl_2_ (0.5µM), 50µl CaCl_2_ (0.1mM), and 500ml PBS). After the addition of 200µl PBSCM to each well, cells were imaged.

#### Fixation of HEK293T cells

To fix the transfected labeled HEK293T cells, 200 µl of 4% formaldehyde in PBS was added to each well of 96-well plate and incubated at room temperature for 30 minutes. The fixative was removed and cells were rinsed twice with PBSCM. To preserve the fluorescence during imaging, 100µl of 90% glycerol in PBS solution was added to each well.

### Microscopy

#### Fluorescence imaging

Confocal fluorescence imaging of cells was performed with a Leica SP2 confocal microscope equipped with an argon laser, a helium/neon laser, and a 63× 1.2-N.A. HCX PL APO water-corrected objective. CFP and HALO-AF488, respectively, were excited on 458 and 488 nm with 50% and 70% power input. CFP and HALO-AF488 were visualized, respectively, with emission spectra of 470-550 and 508-607. The resolution for fluorescence confocal imaging was set to 512×512 pixels. The pinhole size for confocal imaging was set to 133 µm.

For the cells expressing SNAP and HALO domains, on the day of imaging, wells containing SNAP domain were labeled with 5µM SNAP-AF488 for 30 minutes and wells containing HALO domain were labeled with 2 µM of HALO-AF568 for 60 minutes at 37°C. Cells were washed twice with PBSCM. After the addition of 200 µl PBSCM to each well, cells were imaged. Confocal fluorescence imaging of cells was performed with a Leica SP5 confocal microscope equipped with a white light laser (WLL) from Leica Microsystem and an HCX PL APO lambda blue 63.0 × NA 1.20 water objective. SNAP-AF488 and HALO-AF568 were excited respectively on 488 and 578 nm with 70% input power. SNAP-AF488 and HALO-AF568 were visualized, respectively, with emission spectra of 508-607 and 570-700 nm. The resolution for confocal fluorescence imaging was set to 512×512 pixels and the pinhole to 1 airy unit (AU). Confocal images were exported to LAS X Office (Leica Microsystems), and overlay images were obtained with ImageJ.

#### FLIM imaging

For cells expressing CFP and HALO domain, FLIM measurement was performed with a Leica SP2 laser-scanning confocal microscope equipped with a Hamamatsu R3809U-52 photomultiplier detector controlled by a Becker–Hickl time-correlated single-photon counting (TCSPC) control module. Live cells were imaged in 96-well colored-black plates with glass bottom using a 63× 1.2-N.A. water HCX PL APO water-immersion objective with a 405-nm pulsed diode laser as an excitation source (PicoQuant) with input power of 20% for 900s. Resolution for FLIM set to 256×256 pixels and pinhole to 600 µm. Data was exported to SPCImage software version 3.9.7 (Becker–Hickl). The fit method is set to Most Likelihood Estimation (MLE), and the fit model for all analyses is set to mono-exponential. Instrument response function (IRF) for all analyses set to Auto IRF. The threshold was set to 100 counts per pixel to remove background noise. After an initial fit for the whole image was applied, the Region of Interest (ROI) was drawn on the membrane area, and lifetime decay was measured for the chosen ROI. To calculate the E_FRET_ of FRET-enabled AMPARs, the lifetime of the donor was measured in the presence and absence of an acceptor. For this purpose, after measuring τ_D_ and τ_DA_, E_FRET_ was calculated from equation (1) and placed in equation (2) to calculate the *r*.

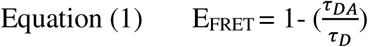

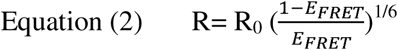

For cells expressing SNAP and HALO domains, FLIM measurement was performed with a Leica SP5 laser-scanning confocal microscope equipped with pulsed WLL SymPhoTime 64, TCSPC module (PicoHarp 300, PicoQuant) and a Hybrid detector (HyD). Resolution for fluorescence lifetime imaging was set to 256×256 pixels. FLIM data was exported to SymPhoTime 64 Confocal TCSPC Data Acquisition and Analysis Software (PicoQuant). After measuring τ_D_ and τ_DA_, E_FRET_ was calculated from equation (1) and by placing E_FRET_ in equation (2) the *r* was calculated.

### Statistical Analysis

Statistical analyses were performed using one-way ANOVA followed by Dunnett’s post hoc tests (GraphPad Prism) to determine the differences between lifetime measurements in HEK293T cells. To compare specific conditions, namely, donor-only lifetime and donor-acceptor lifetime within each experimental group Dunnett’s test was suitable.

