## Supplementary material for "FLIM-FRET Imaging of AMPA Receptors: New Principle for Subtype-Specific Elucidation": SI

**Supplementary Data**

### Material

Dulbecco's Modified Eagle Medium (DMEM), fetal bovine serum (FBS), trypsin, and penicillin/streptomycin (P/S) were purchased from Invitrogen, USA. LipoD293 was purchased from SignaGen, Frederick, MD, USA. All DNA restriction enzymes were from New England Biolabs, USA. PCR enzyme (Phusion Green Hot Start II High-Fidelity PCR Master Mix), Lipofectamine 3000 was purchased from ThermoFisher, USA. Poly-D-Lysine (PDL), formaldehyde, and glycerol were from Sigma-Aldrich. Cell culture dishes were from Sarstedt AG & Co, Germany, and glass bottom plates for microscopy were from Cellvis, USA. The HALO substrates, HALO-AF488 and HALO-AF568 were provided by Matthias M. Herth laboratory (KU, Denmark). SNAP-Surface 488-BG was purchased from Bionordika, Denmark.

**Tabel S1. Primers for PCR amplification of CFP.**

| Primer name | Sequence (5’-3’) |
|---|---|
| FP-GluA1-f | GGTGCCAATTTCCCCGATATCGGAACTAGTGGAGGTATGGTGAGC AAGGGCGAGGAG |
| FP-GluA1-r | GGCATGGACGAGCTGTACAAGGGTGGAACTAGTGGAGATATC ATCCAGATAGGGGGG |
| FP-GluA2-f | GGTGTCTCTTCTAACACCGGTGGAACTAGTGGAGGT ATGGTGAGCAAGGGCGAGG |
| FP-GluA2-r | GGCATGGACGAGCTGTACAAGGGTGGAACTAGTGGAACCGGT CAGATAGGGGGGC |
| FP-GluA3-f | GGATTCCCCAACGATGGAACTAGTGGAGGTATGGTCAGCAAGGGCGAGG |
| FP-GluA3-r | TCCACCTATGCTGATTCCACTAGTTCCACCCTTGTACAGCTCGTCCATGCC |

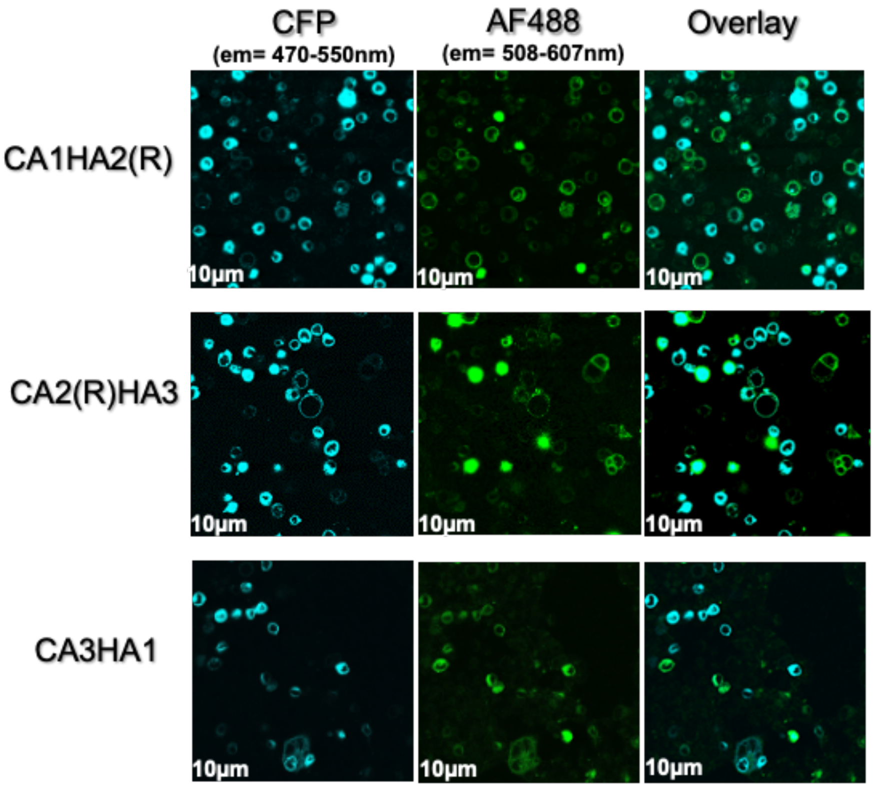


**Figure. S1. Confocal images of transfected HEK293T cells with AMPAR and fluorescence markers** (a) Cells transfected with pmLINK-C-A1-H-A2(R) after labeling individually with AF488 and AF568 are fluorescent. (b) CFP and HALO domain fluorescence signals represent an expression of GluA2(R)$_i$ and GluA3$_i$ (c) CA3HA1 expressed in HEK293T cells.

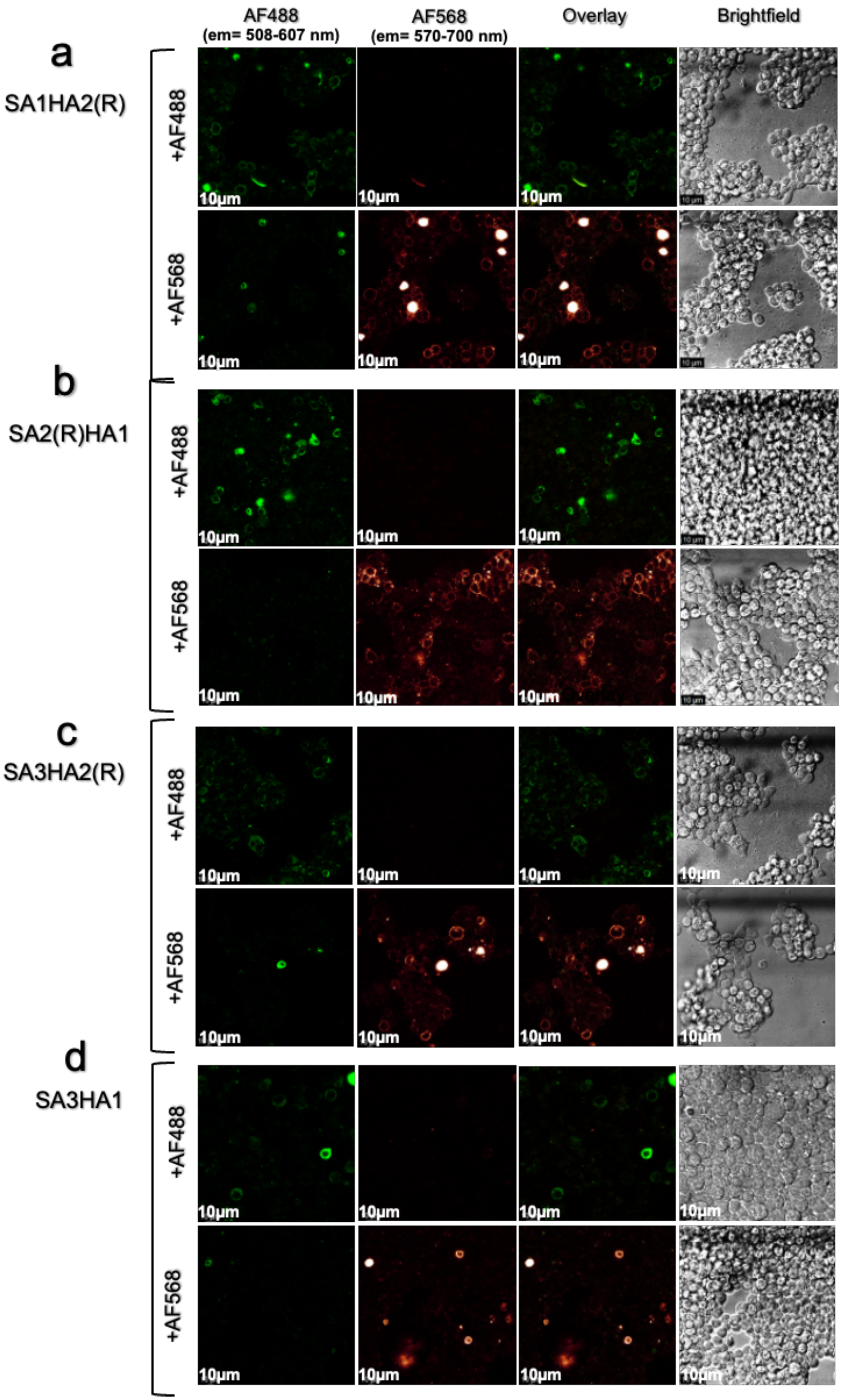
AF488
(em= 508-607 nm)
AF568
(em= 570-700 nm)
Overlay
Brightfield
a
SA1HA2(R)
+AF488
+AF568
b
SA2(R)HA1
+AF488
+AF568
c
SA3HA2(R)
+AF488
+AF568
d
SA3HA1
+AF488
+AF568
10μm

**Figure. S2. Confocal images of transfected HEK293T cells with dual AMPAR and fluorescence tags** Cells transfected with **(a)** pmLINK-S–A1-H-A2(R), **(b)** pmLINK-H-A1-S-A2(R), **(c)** pmLINK-H-A2(R)-S-A3, and **(d)** pmLINK-S-A3-H-A1 after labeling with AF488 and AF568 are fluorescence.
